# Representation of *k*-mer sets using spectrum-preserving string sets

**DOI:** 10.1101/2020.01.07.896928

**Authors:** Amatur Rahman, Paul Medvedev

## Abstract

Given the popularity and elegance of *k*-mer based tools, finding a space-efficient way to represent a set of *k*-mers is important for improving the scalability of bioinformatics analyses. One popular approach is to convert the set of *k*-mers into the more compact set of unitigs. We generalize this approach and formulate it as the problem of finding a smallest spectrum-preserving string set (SPSS) representation. We show that this problem is equivalent to finding a smallest path cover in a compacted de Bruijn graph. Using this reduction, we prove a lower bound on the size of the optimal SPSS and propose a greedy method called UST that results in a smaller representation than unitigs and is nearly optimal with respect to our lower bound. We demonstrate the usefulness of the SPSS formulation with two applications of UST. The first one is a compression algorithm, UST-Compress, which we show can store a set of *k*-mers using an order-of-magnitude less disk space than other lossless compression tools. The second one is an exact static *k*-mer membership index, UST-FM, which we show improves index size by 10-44% compared to other state-of-the-art low memory indices. Our tool is publicly available at: https://github.com/medvedevgroup/UST/.

## 1 Introduction

Algorithms based on *k*-mers are now amongst the the top performing tools for many bioinformatics analyses. Instead of working directly with reads or alignments, these tools work with the set of *k*-mer substrings present in the data, often relying on specialized data structures for representing sets of *k*-mers (for a survey, see [1]). Since modern sequencing datasets are huge, the space used by such data structures is a bottleneck when attempting to scale up to large databases. For example, as part of our group’s work on building indices for RNA-seq data, we are storing gzipped *k*-mer set files from about 2,500 experiments [2]. Though this is only a fraction of experiments in the SRA, it already consumes 6 TB of space. For these and other applications, the development of space-efficient representations of *k*-mer sets can improve scalability and enable novel biological discoveries.

Conway and Bromage [3] showed that at least 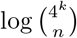 bits are needed to losslessly store a set of *n k*-mers, in the worst case. However, a set of *k*-mers generated from a sequencing experiment typically exhibits the spectrum-like property [1] and contains a lot of redundant information. Therefore, in practice, most data structures can substantially improve on that bound [4].

A common way to reduce the redundancy in a *k*-mer set *K* is to convert it into a set of *maximal unitigs*. A unitig is a non-branching path in the de Bruijn graph, a graph whose nodes are the *k*-mers of *K* and edges are the overlaps between *k*-mers. A unitig *u* can be written as a string *spell*(*u*) of length *|u|* + *k −* 1, such that the *k*-mers of *u* are exactly the *k*-mer substrings of *spell*(*u*). For example, the unitig (*AAC, ACG, CGT*) is spelled as *AACGT*. This gives a way to represent *|u| k*-mers using *|u|* + *k −* 1 characters, instead of *k|u|* characters used by a naive approach. When unitigs are long, as they are in real data, the space savings are significant. The idea can be extended to store the whole set *K*, because the set of maximal unitigs *U* forms a decomposition of *K*, and, therefore, has the nice property that *x ∈ K* iff *x* is a substring of *spell*(*u*), for some *u ∈ U*.

The maximal unitigs *U* can be computed efficiently [5–7] and combined with an auxiliary index to obtain a membership data structure (i.e. one that can efficiently determine if a *k*-mer belongs to *K* or not). In particular, Unitigs-FM [4] and deGSM [7] uses the FM-index as the auxiliary index, Pufferfish [8] and BLight [9] uses a minimum perfect hash function, and Bifrost [10] uses a minimizer hash table. Alternatively, *U* can be compressed to obtain a compressed disk representation of *K*, albeit without efficient support for membership queries prior to decompression.

While unitigs conveniently fit the needs of those applications, we observe in this paper that they are not necessarily the best that can be done. Concretely, we claim that what makes *U* useful in these scenarios is that they are a type of *spectrum-preserving string set (SPSS) representation* of *K*, which we define to be a set of strings *X* such that a *k*-mer is in *K* iff it is a substring of a string in *X*. (This is in contrast to the way unitigs are used in assembly, where it is crucial that they are not chimeric [11].) The *weight* of *X* is the number of characters it contains. In this paper, we explore the idea of low weight representations and their applicability. In particular, are there representations with a smaller weight than *U* that can be efficiently computed? What is the lowest weight that is achievable by a representation? Can such representations seamlessly replace unitig representations in downstream applications, and can they improve space performance?

In this paper, we show that the problem of finding a minimum weight SPSS representation is equivalent to finding a smallest path cover in a compacted de Bruijn graph (Section 3). We use the reduction to give a lower bound on the weight which could be achieved by any SPSS representation (Section 4), and we give an efficient greedy algorithm UST to find a representation that improves on *U* (Section 5) and is empirically near-optimal. We demonstrate the usefulness of our representation using two applications (Section 6). One, we combine it with an FM-index into a membership data structure called UST-FM, and, two, we combine it with a general compression algorithm to give a compression algorithm called UST-Compress. Both applications result in a substantial space decrease over state-of-the-art (Section 7), demonstrating the usefulness of SPSS representations. Our software is freely available at https://github.com/medvedevgroup/UST/.

### 1.1 Related work

The idea of using a SPSS for a membership index was previously independently described in a PhD thesis [12] and questions similar to the ones in our paper are simultaneously and independently studied in [13]. The idea of greedily gluing unitigs (as UST does) has previously appeared in read compression [14], where contigs greedily constructed from the reads and the reads were stored as alignments to these contigs. The idea also appeared in the context of sequence assembly, where a greedy traversal of an assembly graph was used as an intermediate step during assembly [15, 16].

The compression of *k*-mer sets has not been extensively studied, except in the context of how *k*-mer counters store their output [17–20]. DSK [18] uses an HDF5-based encoding, KMC3 [17] combines a dense storage of prefixes with a sparse storage of suffixes, and Squeakr [20] uses a counting quotient filter [21]. The compression of read data, on the other hand, stored in either unaligned or aligned formats, has received a lot of attention [22–24]. In the scenario where the *k*-mer set to be compressed was originally generated from FASTA files by a *k*-mer counter, an alternate to *k*-mer compression is to compress the original FASTA file and use a *k*-mer counter as part of the decompression to extract the *k*-mers on the fly. This approach is unsatisfactory because 1) as we show in this paper, it takes substantially more space than direct *k*-mer compression, 2) *k*-mer counting on the fly adds significant time and memory to the decompression process, and 3) there are applications where the *k*-mer set cannot be reproduced by simply counting *k*-mers in a FASTA file, e.g. when it is a product of a multi-sample error correction algorithm [25]. Furthermore, there are applications where the *k*-mer set is not related to sequence read data at all, e.g. a universal hitting set [26], a chromosome-specific reference dictionary [27], or a winnowed min-hash sketch (for example as in [28], or see [29, 30] for a survey).

Membership data structures for *k*-mer sets were surveyed in a recent paper [1]. In addition to the unitig-based approaches already mentioned, other exact representations include succinct de Bruijn graphs (referred to as BOSS [31]) and their variations [32, 33], dynamic de Bruijn graphs [34, 35], and Bloom filter tries [36]. Some data structures are non-static, i.e. they provide the ability to insert and/or delete *k*-mers. However, such operations are not needed in many read-only applications, where the cost of supporting them can be avoided. Membership data structures can be extended to associate additional information with each *k*-mer, for instance an abundance count (e.g. deBGR [37]) or a color class (for a short overview, see [1]).

## 2 Definitions

*Strings:* In this paper, we assume all strings are over the alphabet *Σ* = *{A, C, G, T}*. The *length* of string *x* is denoted by *|x|*. A string of length *k* is called a *k-mer*. For a set of strings *S*, *weight*(*S*) = ^),^*_x∈S_ |x|* denotes the total count of characters. We write *x*[*i..j*] to denote the substring of *x* from the *i*^th^ to the *j*^th^ character, inclusive. We define *suf_k_* (*x*) (respectively, *pre_k_* (*x*)) to be the last (respectively, first) *k* characters of *x*. For *x* and *y* with *suf_k−_*_1_(*x*) = *pre_k−_*_1_(*y*), we define *gluing x* and *y* as *x 8 y* = *x · y*[*k..|y|*]. For *s ∈ {*0, 1*}*, we define *orient*(*x, s*) to be *x* if *s* = 0 and to be the reverse complement of *x* if *s* = 1. A string *x* is *canonical* if *x* is the lexicographically smaller of *x* and its reverse complement. To *canonize x* is to replace it by its canonical version (i.e. min*_i_*(*orient*(*x, i*))). We say that *x*_0_ and *x*_1_ have a (*s*_0_*, s*_1_)*-oriented-overlap* if *suf_k−_*_1_(*orient*(*x*_0_, 1 *−s*_0_) = *pre_k−_*_1_(*orient*(*x*_1_*, s*_1_)). Intuitively, such an overlap exists between two strings if we can orient them in such a way that they are glueable. We define the *k-spectrum sp^k^* (*x*) as the multi-set of all canonized *k*-mer substrings of *x*. The *k*-spectrum for a set of strings *S* is defined as *sp^k^* (*S*) = ⋃ *_x ∈ S_ sp^k^* (*x*).

*Bidirected graphs:* A *bidirected graph G* is a pair (*V, E*) where the set *V* are called vertices and *E* is a set of edges. An edge *e* is a 4-tuple (*u*_0_*, s*_0_*, u*_1_*, s*_1_), where *u_i_ ∈ V* and *s_i_ ∈ {*0, 1*}*, for *i ∈ {*0, 1*}*. Intuitively, every vertex has two sides, and an edge connects to a side of a vertex. Note that there can be multiple edges between two vertices, but only one edge once the sides are fixed. An edge is a *loop* if *u*_0_ = *u*_1_. Given a non-loop edge *e* that is incident to a vertex *u*, we denote *side*(*u, e*) as the side of *u* to which it is incident. We say that a vertex *u* is *isolated* if it has no edge incident to it and is a *dead-end* if it has exactly one side to which no edges are incident. Define *n_dead_* and *n_iso_* as the number of dead-end and isolated vertices, respectively. A sequence *w* = (*u*_0_*, e*_1_*, u*_1_*, …, e_n_, u_n_*) is a *walk* if for all 1 *≤ i ≤ n*, *e_i_* is incident to *u_i−_*_1_ and to *u_i_*, and for all 1 *≤ i ≤ n −* 1, *side*(*u_i_, e_i_*) = 1 *− side*(*u_i_, e_i_*_+1_). Vertices *u*_1_*, …, u_n−_*_1_ are called *internal* and *u*_0_ and *u_n_* are called *endpoints*. A walk can also be a single vertex, in which case it is considered to have no internal vertex and one endpoint. A *path cover W* of *G* is a set of walks such that every vertex is in exactly one walk in *W* and no walk visits a vertex more than once.

*Bidirected DNA graphs:* A *bidirected DNA graph* is a bidirected graph *G* where every vertex *u* has a string label *lab*(*u*), and for every edge *e* = (*u*_0_*, s*_0_*, u*_1_*, s*_1_), there is a (*s*_0_*, s*_1_)-oriented-overlap between *lab*(*u*_0_) and *lab*(*u*_1_). *G* is said to be *overlap-closed* if there is an edge for every such overlap. Let *w* = (*u*_0_*, e*_1_*, u*_1_*, …, e_n_, u_n_*) be a walk. Define *x*_0_ = *orient*(*lab*(*u*_0_), 1 *− side*(*u*_0_*, e*_1_)) and, for 1 *≤ i ≤ n*, *x_i_* = *orient*(*lab*(*u_i_*)*, side*(*u_i_, e_i−_*_1_)). The *spelling* of a walk is defined as *spell*(*w*) = *x*_0_ *8 · · · 8 x_n_*. (The fact that the *x_i_*’s are glueable in this way can be derived from definitions.) If *W* is a set of walks, then define *spell*(*W*) = ⋃ *_w ∈ W_ spell*(*w*).

*De Bruijn graphs:* Let *K* be a set of canonical *k*-mers. The node-centric *bidirected de Bruijn graph*, denoted by *dBG*(*K*), is the overlap-closed bidirected DNA graph where the vertices and their labels correspond to *K*. Figure 1A shows an example. In this paper, we will assume that *dBG*(*K*) is not just a single cycle; such a case is easy to handle in practice but is a space-consuming corner-case in all the analyses. A walk in *dbG*(*K*) is a *unitig* if all its vertices have in- and out-degrees of 1, except that the first vertex can have any in-degree and the last vertex can have any out-degree.

**Fig. 1:**
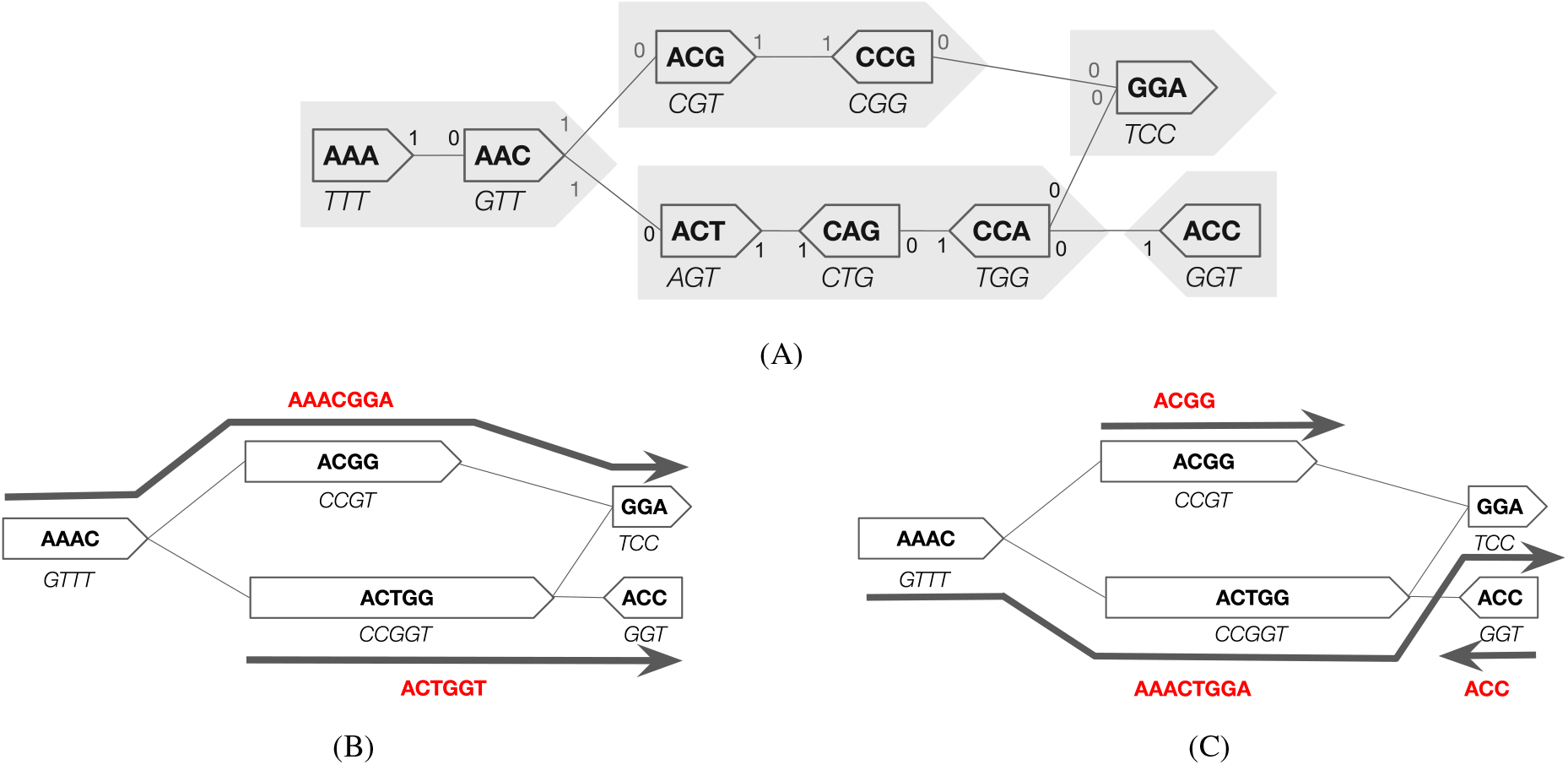
*(A)* An example of a de Bruijn graph for a set *K* with 9 3-mers. The 0 side of a vertex is drawn flat and the 1 side pointy. The text in each vertex is its label, i.e. what is spelled by a walk going in the direction of the pointy end. The string below the vertex is the reverse complement of its label, which is what is spelled by a walk going in the opposite direction. The maximal unitigs are shown by filled in gray arrows. *(B)* The compacted de Bruijn graph for the same set *K*. Each vertex corresponds to a maximal unitig in the top graph. Each vertex’s label corresponds to the spelling of the corresponding unitig and is shown inside the vertex; the reverse complement of the label is written below in italics. One possible path cover is five walks, each corresponding to a single vertex; the spelling of this cover is *{AAAC, ACGG, ACTGG, GGA, ACC}*, which is the unitig SPSS representation of *K*. A better path cover of size 2 that could potentially be found by our UST algorithm is shown. It corresponds to SPSS representation *{AAACGGA, ACTGGT}*. It is easy to verify that this path cover has minimum size, and, by Theorem 1, the corresponding representation has minimum weight (13). *(C)* Another path cover that could potentially be found by UST. It has size 3 and is suboptimal.

A single vertex is also a unitig. A unitig is *maximal* if it is not a sub-walk of another unitig. It was shown in [5] that if *dBG*(*K*) is not a cycle, then a unitig cannot visit a vertex more than once, and the set of maximal unitigs forms a unique decomposition of the vertices in *dBG*(*K*) into non-overlapping walks. The bidirected *compacted de Bruijn graph* of *K*, denoted by *cdBG*(*K*), is the overlap-closed bidirected DNA graph where the vertices are the maximal unitigs of *dBG*(*K*), and the labels of the vertices are the spellings of the unitigs. Figure 1B shows an example.

## 3 Equivalence of SPSS representations and path covers

For this section, we fix *K* to be a canonical set of *k*-mers. A set of strings *X* is said to be a *spectrum-preserving string set (SPSS) representation* of *K* iff their *k*-spectrums are equal and each string in *X* is of length ≥ *k*. For brevity, we say *X represents K*. Note that because in our definitions *K* is a set (i.e. no duplicates) and the *k*-spectrum is a multi-set, this effectively restricts *X* to not contain duplicate *k*-mers. See Figure 1BC for examples. In this paper, we consider the problem of finding a minimum weight SPSS representation of *K*. In this section, we will show that it is equivalent to the problem of finding a smallest path cover of *cdBG*(*K*), in the following sense:

### Theorem 1.

*Let X*^*opt*^ *be a minimum weight SPSS representation of K. Let W* ^*opt*^ *be the smallest path cover on cdBG(K). Then, weight*(*X*^*opt*^) = |*K*| + |*W*^*opt*^|(*k* − 1).

First, we show that the weight of a SPSS representation is a linear increasing function of its size (i.e. the number of strings it contains) and, hence, finding a SPSS representation of minimum weight is equivalent to finding one of minimum size.

### Lemma 1.

*Let X be a SPSS representing K. Then, weight*(*X*) = |*K*| + |*X*|(*k* − 1).

*Proof.* Every string *x* of length ≥ *k* cLontains |*x*| − *k* + 1 *k*-mers. *X* has |*K*| *k*-mers, since *X* and *K* have the same *k*-spectrum. Combining these, |*K*| = ∑_*x*∈*X*_(|*x*| − *k* + 1) = *weight*(*X*) − |*X*|(*k* − 1).

The intuition behind Theorem 1 is that there is a natural size-preserving bijection between path covers of *dBG*(*K*) and SPSS reprsentations of *K*. Since it is more efficient to work with compacted de Bruijn graphs, we would like this to hold for *cdBG*(*K*) as well. However, the path covers with an endpoint at an internal vertex of a unitig in *dBG*(*K*) do not project onto *cdBG*(*K*). Nevertheless, this is not an issue because such path covers are necessarily non-optimal.

### Lemma 2.

*Let W be a path cover of cdBG(K). Then spell(W) represents K*.

*Proof.* By construction, all strings in *spell*(*W*) are at least *k*-long, so we only need to show that the spectrum of *spell*(*W*) is *K*. Let *W*^0^ be the path cover with every vertex as its own walk. We can view *W* as being constructed from *W*^0^ by repeatedly taking a pair of walks that share endpoints and joining them together. We prove the Lemma by induction. For the base case, *spell*(*W*^0^) are the unitigs of *dBG*(*K*), which, by definition, have the same spectrum as *K* [5]. Now let *W*^*i*^ be the path cover after *i* walk-joins. Then *W*^*i*+1^ is the result of joining some two walks *w* and *w*′ into *w*′′. Observe that joining walks preserves the *k*-spectrum of their spellings, i.e *sp*^*k*^(*spell*(*w*)) ⋃ *sp*^*k*^(*spell*(*w*′)) = *sp*^*k*^(*spell*(*w*′′)). Combining with the inductive hypothesis for *W*^*i*^, *sp*^*k*^(*spell*(*W*^*i*^)) = *sp*^*k*^(*spell*(*W*^*i*+1^)).

### Lemma 3.

*Let X be a smallest SPSS representation of K. Then there exists a path cover W of cdBG(K) with* |*W* | = |*X*|.

*Proof.* Let *X* = {*x*_1_, …, *x*_*m*_}. Every string *x*_*i*_ is spelled by a walk *w*′_*i*_ in *dBG*(*K*), visiting the sequence of its canonized constituent *k*-mers. Since *X* is spectrum preserving with respect to *K*, it contains every *k*-mer in *K* exactly once; therefore, {*w*′_1_, …, *w*′_*m*_} is a path cover of *dbG*(*K*).

Since *X* has the smallest number of strings, the endpoints of *w*′_*i*_ cannot be on internal vertices of unitigs, otherwise there would exist another string *x*_*j*_ that could be glued with *x*_*i*_ to form a smaller SPSS representing *K*. Therefore, there exists a corresponding walk *w*_*i*_ in *cdBG*(*K*) such that *spell*(*w*_*i*_) = *spell*(*w*′_*i*_) = *x*_*i*_. Hence, the set of walks *W* = {*w*_1_, …, *w*_*m*_} is a path cover of *cdBG*(*K*).

Now we can prove Theorem 1.

*Proof.* By Lemma 1, *X*^opt^ has minimum size and, hence, by Lemma 3, there exists a path cover *W* with |*W*| = |*X*^opt^7. By the optimality of *W*^opt^, |*W*^opt^| ≤ |*W* ≤ |*X*^opt^. Next, by Lemma 2, *spell*(*W*^opt^) represents *K* and, by definition, |*spell*(*W*^opt^) = |*W*^opt^|. Since *X*^opt^ has minimum size, |*X*^opt^| ≤ |*spell*(*W*^opt^)| = |*W*^opt^|. This proves |*X*^opt^| = |*W*^opt^|. Lemma 1 then implies the Theorem.

## 4 Lower bound on the weight of a SPSS representation

In this section, we will prove a lower bound on the size of a path cover of a bidirected graph, which, by Theorem 1, gives a lower bound on the weight of any SPSS representation. Finding the minimum size of a path cover in general directed graphs is NP-hard, since a directed graph has a Hamiltonian path if and only if it has a path cover of size 1. However, we do not know the complexity of the problem when restricted to compacted de Bruijn graphs of *k*-mer sets. The minimum size of a path cover is known to be bounded from above by the maximum size of an independent set (at least for directed graphs [38]); however, finding a maximum independent set is itself NP-hard. We therefore take a different approach.

For this section, let *G* = (*V, E*) be a bidirected graph without loops and let *W* be a path cover. A *vertex-side* is a pair (*u, su*), where *u ∈ V* and *su ∈ {*0, 1*}*. For a non-isolated vertex *u*, we say (*u, su*) is a *dead-side* if there are no edges incident to (*u,* 1*−su*). Note that the number of dead-sides is by definition the number of dead-end vertices. Consider a walk (*v*_0_*, e*_1_*, …, e_n_, v_n_*) with *n ≥* 1. Denote its *endpoint-sides* as (*v*_0_*, side*(*v*_0_*, e*_1_)) and (*v_n_, side*(*v_n_, e_n_*)). If a walk contains just one vertex (*v*_0_), then denote its *endpoint-sides* as (*v*_0_, 0) and (*v*_0_, 1).

We observe that every walk in a path cover must have two unique endpoint-sides. Our strategy is to give a lower bound on the number of endpoint-sides, thereby giving a lower bound on the size of a path cover. We know, for instance, that dead-sides must be endpoint-sides and we also know that the sides of an isolated vertex must be endpoint-sides. For other cases, we cannot predict exactly the endpoint-sides, but we can create disjoint sets of vertex-sides (which we call special neighborhoods) such that, for each set, we can guarantee that all but one of its vertex-sides are endpoint-sides. Formally, for a vertex-side (*u, su*), its *special neighborhood B_u,su_* is the set of vertex-sides (*v, sv*) such that there exists and edge between (*u, su*) and (*v,* 1 *− sv*) and it is the only edge incident on (*v,* 1 *− sv*). A vertex-side which belongs to a special neighborhood is called a *special-side*. Figure 2 shows an example. Our key lemma is that all but one member of a special neighborhood must be an endpoint-side:

**Fig. 2:**
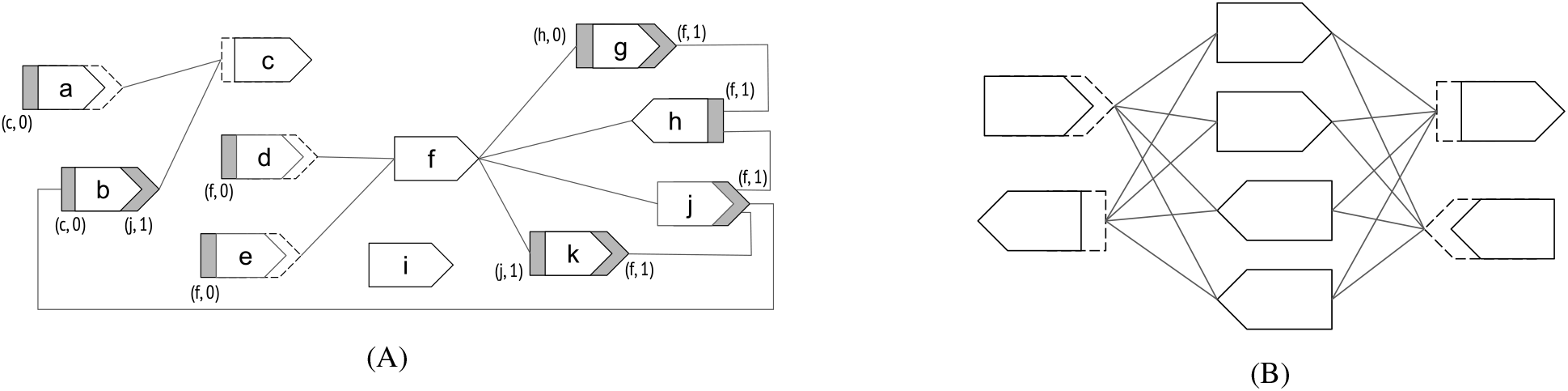
(A) An example compacted de Bruijn graph (labels not shown), with a distinct ID for each vertex shown inside the vertex. The dashed hollow sides of the vertices are dead sides and the solid gray sides are special-sides. Each special-side is additionally labeled with the vertex-side to whose special neighborhoods it belongs. For example, the special neighborhood of vertex-side (c, 0) contains two vertex-sides, namely the blunt gray sides of vertices a and b, corresponding to *|Bc,*0*|* = 2. In this example, *ndead* = 4, *nsp* = 6, and *niso* = 1. By Theorem 2, the minimum size of a path cover is 6, and one can indeed find a path cover of this size in the graph. (B) In this example, *ndead* = 4, *nsp* = *niso* = 0, resulting in a lower bound of 2 on the size of a path cover. However, a quick inspection tells us that the the optimal size of a path cover is 4. This shows that our lower bound is not theoretically tight.

### Lemma 4.

*For a vertex-side* (*u, su*), *there must be at least* |*B*_*u,su*_| − 1 *endpoint-sides of W in B*_*u,su*_.

*Proof.* Assume without loss of generality that |*B*_*u,su*_| > 1, since the lemma is vacuous otherwise. Let (*v, sv*) ∈ *B*_*u,su*_ be a vertex-side that is not an endpoint side in *W*, and let *w*_*v*_ be the walk containing *v*. Since, in particular, (*v, sv*) is not an endpoint-side of *w*_*v*_, then *w*_*v*_ must contain an edge incident to (*v*, 1 − *sv*). By definition of special neighborhood, the only such edge is incident to (*u, su*). By definition of a path cover, there can only be one walk in *W* that contains an edge incident to (*u, su*) and it can contain only one such edge. Hence, there can only be one (*v, sv*) ∈ *B*_*u,su*_ that is not an endpoint-side.

Next, we show that the special neighborhoods are disjoint, and we can therefore define *n*_*sp*_ = ∑_*u*∈*V,su*∈{0,1}_ max(0, |*B*_*u,su*_| − 1) as a lower bound on the number of special-sides that are endpoint-sides:

### Lemma 5.

*There are at least n*_*sp*_ *special-sides that are endpoint-sides of W*.

*Proof.* We claim that *B*_*u,su*_ ∩ *B*_*v,sv*_ = ϕ for all (*u, su*) ≠ (*v, su*). Let (*w, sw*) ∈ *B*_*u,su*_ ∩ *B*_*v,sv*_. By definition of *B*_*u,su*_, the only edge touching (*w*, 1−*sw*) is incident to (*u, su*). Similarly, by definition of *B*_*v*_, the only edge touching (*w*, 1−*sw*) is incident to (*v, sv*). Hence, (*u, su*) = (*v, sv*). The Lemma follows by applying Lemma 4 to each *B*_*u,su*_ and summing the result.

Finally, we are ready to prove our lower bound on the size of path cover.

### Theorem 2.

|*W*| ≥ [(*n*_*dead*_ + *n*_*sp*_)/2] + *n*_*iso*_.

*Proof.* Define a walk as *isolated* if it has only one vertex and that vertex is isolated. There are exactly *n*_*iso*_ isolated walks in *W*. Next, dead-sides are trivially endpoint-sides of a non-isolated walk in *W*. By Lemma 5, so are at least *n*_*sp*_ of the special-sides. Since the set of dead-sides and the set of special-sides are, by their definition, disjoint, the number of distinct endpoint-sides of non-isolated walks is at least *n*_*dead*_ + *n*_*sp*_. Since every walk in a path cover must have exactly two distinct endpoint-sides, there must be at least [(*n*_*dead*_ + *n*_*sp*_)/2] non-isolated walks.

By applying Theorem 1 to Theorem 2 and observing that loops do not effect path covers, we get a lower bound on the minimum weight of any SPSS representation:

### Corollary 1.

*Let K be a set of canonical k-mers and let X*^*opt*^ *be its minimum weight SPSS representation. Then, weight*(*X*^*opt*^) ≥ |*K*| + (*k* − 1) ([(*n*_*dead*_ + *n*_*sp*_)/2] + *n*_*iso*_), *where n*_*dead*_, *n*_*iso*_, *and n*_*sp*_ *are defined with respect to the graph obtained by removing loops from cdBG*(*K*).

We note that the lower bound is not tight, as in the example of Figure 2B; it can likely be improved by accounting for higher-order relationships in *G*. However, the empirical gap between our lower bound and algorithm is so small (Section 7) that we did not pursue this direction.

## 5 The UST algorithm for computing a SPSS representation

In this section, we describe our algorithm called UST (UNITIG-STITCH) for computing a SPSS representation of a set of *k*-mers *K*. We first use the Bcalm2 tool [5] to construct *cdBG*(*K*), then find a path cover *W* of *cdBG*(*K*), and then output *spell*(*W*), which by Lemma 2 is a SPSS representation of *K*.

UST constructs a path cover *W* by greedily exploring the vertices, with each vertex explored exactly once. We maintain the invariant that *W* is a path cover over all the vertices explored up to that point, and that the currently explored vertex is an endpoint of a walk in *W*. To start, we pick an arbitrary vertex *u*, add a walk consisting of only *u* to *W*, and start an *exploration* from *u*.

An exploration from *u* works as follows. First, we mark *u* as explored. Let *w_u_* be the walk in *W* that contains *u* as an endpoint, and let *su* be the endpoint-side of *u* in *w_u_*. We then search for an edge *e* = (*u,* 1 *− su, v, sv*), for some *v* and *sv*. If we find such an edge and *v* has not been explored, then we extend *w_u_* with *e* and start a new exploration from *v*. If *v* has been explored and is an endpoint vertex of a walk *w_v_* in *W*, then we merge *w_u_* and *w_v_* together if the orientations allow (i.e. if 1 *− sv* is the side at which *w_v_* is incident to *v*) and start a new exploration from an arbitrary unexplored vertex. In all other cases (i.e. if *e* is not found, if the orientations do not allow merging *w_v_* with *w_u_*, or if *v* in internal vertex in *w_v_*), we start a new exploration from an arbitrary unexplored vertex. The algorithm terminates once all the vertices have been explored. It follows directly via the loop invariant that the algorithm finds a path cover, though we omit an explicit proof.

In our implementation, we do not store the walks *W* explicitly but rather just store a walk ID at every vertex along with some associated information. This makes the algorithm run-time and memory linear in the number of vertices and the number of edges, except for the possibility of needing to merge walks (i.e. merging of *w_u_* and *w_v_*). But we implement these operations using a union-find data structure, making the total time near-linear.

We note that UST’s path cover depends on the arbitrary choices of which vertex to explore. Figure 1C gives an example where this leads to sub-optimal results. However, our results indicate that UST cannot be significantly improved in practice, at least for the datasets we consider (Section 7).

## 6 Applications

We apply UST to solve two problems. First, we use it to construct a *compression* algorithm UST-Compress. UST-Compress supports only compression and decompression and not membership and is intended to reduce disk space. We take *K* as input (in the binary output format of either DSK [18] or Jellyfish [19]), run UST on *K*, and then compress the resulting SPSS using a generic nucleotide compressor MFC [39]. UST-Compress can also be run in a mode that takes as input a count associated with each *k*-mer. In this mode, it outputs a list of counts in the order of their respective *k*-mers in the output SPSS representation (this is a trivial modification to UST). This list is then compressed using the generic LZMA compression algorithm. Note that we use MFC and LZMA due to their superior compression ratios, but other compressors could be substituted. To decompress, we simply run the MFC or LZMA decompressing algorithm.

Second, we use UST to construct an *exact static membership data structure* UST-FM. Given *K*, we first run UST on *K*, and then construct an FM-index [40] (as implemented in [41]) on top of the resulting SPSS representation. The FM-index then supports membership queries. In comparison to hash-based approaches, the FM-index does not support insertion or deletion; on the other hand, it allows membership queries of strings shorter than *k*.

## 7 Empirical results

We use different types of publicly available sequencing data because each type may result in a de Bruijn graph with different properties and may inherently be more or less compressible. Our datasets include human, bacterial, and fish samples; they also include genomic, metagenomic, and RNA-seq data (Table S1). Each dataset was *k*-mer counted using DSK [18], using *k* = 31 with singleton *k*-mers removed. While these are not the optimal values for each of the respective applications, it allows us to have a uniform comparison across datasets. In addition, we *k*-mer count one of the datasets with *k* = 61, removing singletons, in order to study the effect of *k*-mer size. All our experiments were run on a server with a Intel(R) Xeon(R) CPU E5-2683 v4 @ 2.10GHz with 64 cores and 512 GB of memory. All tested algorithms were verified for correctness in all datasets. Table S2 shows the version numbers of all tools tested and further reproducibility details are available at https://github.com/medvedevgroup/UST/tree/master/experiments.

**Table 1:** Comparison of different string set representations and the SPSS lower bound. The second column shows *|K|*. For a representation *X*, the number of strings is *|X|* and the number of nucleotides per distinct *k*-mer is *weight*(*X*)*/|K|*. Unitigs were computed using BCALM2.

| Dataset | # distinct $k$-mers | SPSS lower bound | | UST | | unitigs | |
| --- | --- | --- | --- | --- | --- | --- | --- |
| | | # strings | nt / $k$-mer | # strings | nt / $k$-mer | # strings | nt / $k$-mer |
| Zebrafish RNA-seq | 124,740,993 | 3,979,856 | 1.96 | 4,174,867 | 2.00 | 7,775,719 | 2.87 |
| Human RNA-seq | 101,017,526 | 3,924,803 | 2.17 | 4,132,115 | 2.23 | 7,665,682 | 3.28 |
| Human chromosome 14 | 99,941,572 | 2,235,267 | 1.67 | 2,386,324 | 1.72 | 4,871,245 | 2.46 |
| Whole human genome | 391,766,120 | 13,964,825 | 2.07 | 14,423,449 | 2.10 | 19,581,835 | 2.50 |
| Human gut metagenome | 103,814,001 | 1,517,107 | 1.34 | 1,522,139 | 1.34 | 2,187,669 | 1.49 |
| Human RNA-seq ($k=61$) | 75,013,109 | 2,651,729 | 3.12 | 2,713,825 | 3.17 | 4,371,173 | 4.50 |

**Table 2:**
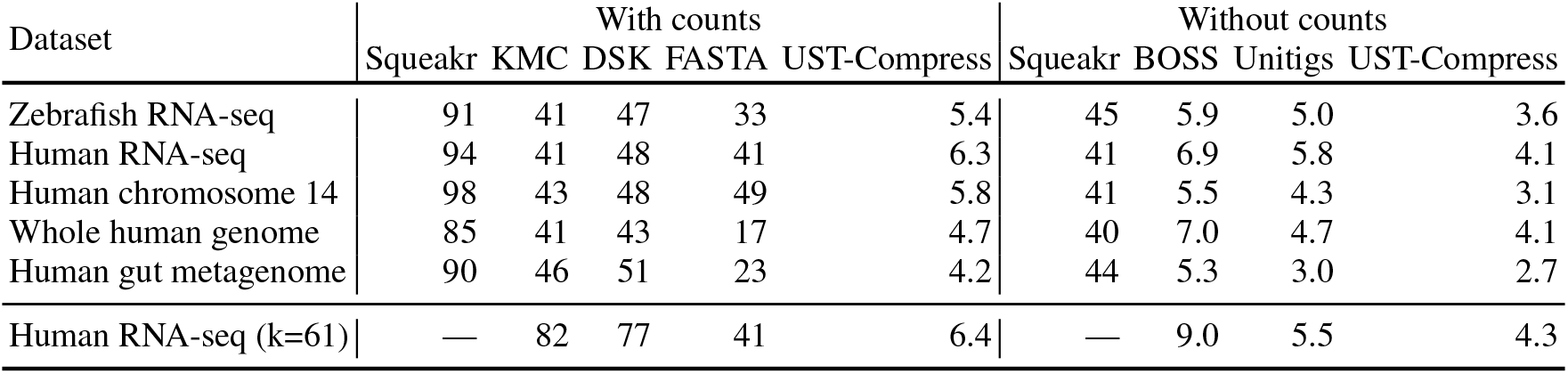
Space usage of UST-Compress and others. We show the average number of bits per distinct *k*-mer in the dataset. All files are compressed with MFC or LZMA, in addition to the tool shown in the column name. Squeakr-exact’s implementation is limited to *k <* 32 [20] and so it could not be run for *k* = 61.

| Dataset | With counts |  |  |  |  | Without counts |  |  |  |
| --- | --- | --- | --- | --- | --- | --- | --- | --- | --- |
|  | Squeakr | KMC | DSK | FASTA | UST-Compress | Squeakr | BOSS | Unitigs | UST-Compress |
| Zebrafish RNA-seq | 91 | 41 | 47 | 33 | 5.4 | 45 | 5.9 | 5.0 | 3.6 |
| Human RNA-seq | 94 | 41 | 48 | 41 | 6.3 | 41 | 6.9 | 5.8 | 4.1 |
| Human chromosome 14 | 98 | 43 | 48 | 49 | 5.8 | 41 | 5.5 | 4.3 | 3.1 |
| Whole human genome | 85 | 41 | 43 | 17 | 4.7 | 40 | 7.0 | 4.7 | 4.1 |
| Human gut metagenome | 90 | 46 | 51 | 23 | 4.2 | 44 | 5.3 | 3.0 | 2.7 |
| Human RNA-seq (k=61) | — | 82 | 77 | 41 | 6.4 | — | 9.0 | 5.5 | 4.3 |

### 7.1 Evaluation of the UST representation

We compare our UST representation against the unitig representation as well as against the SPSS lower bound of Corollary 1 (Table 1, with a deeper breakdown in Table S3). UST reduces the number of nucleotides (i.e. weight) compared to the unitigs by 10-32%, depending on the dataset. The number of nucleotides obtained is always within 3% of the SPSS lower bound; in fact, when considering the gap between the unitig representation and the lower bound, UST closes 92-99% of that gap. These results indicate that our greedy algorithm is a nearly optimal SPSS representation, on these datasets. They also indicate that the lower bound of Corollary 1, while not theoretically tight, is nearly tight on the type of real data captured by our experiments.

**Table 3:** Time and peak memory usage of UST-Compress (without counts) and others during compression. For BOSS and unitigs, the times are separated according to the two steps of compression: running the core algorithm (Cosmo and bcalm2) followed by the generic compressor (respectively, LZMA and MFC). For UST-Compress, the first step is exactly the same as for unitigs (Bcalm2), so the column is not repeated.

| Dataset | Time (minutes) |  |  |  |  |  |  |  |  | Peak memory (GB) |  |  |
| --- | --- | --- | --- | --- | --- | --- | --- | --- | --- | --- | --- | --- |
|  | BOSS |  |  | unitigs |  |  | UST-Compress |  |  | BOSS | unitigs | UST-Compress |
|  | Cosmo | LZMA | Total | bcalm2 | MFC | Total | UST | MFC | Total |  |  |  |
| Zebrafish RNA-seq | 6.3 | 0.7 | 7.0 | 3.0 | 1.5 | 4.4 | 1.5 | 0.9 | 5.3 | 4.0 | 3.1 | 3.1 |
| Human RNA-seq | 4.0 | 0.8 | 4.8 | 4.7 | 1.3 | 5.9 | 1.6 | 0.8 | 7.1 | 3.6 | 3.4 | 3.4 |
| Human chromosome 14 | 4.9 | 0.5 | 5.4 | 2.1 | 1.0 | 3.1 | 1.1 | 0.7 | 3.9 | 4.2 | 3.4 | 3.4 |
| Whole human genome | 17.3 | 3.0 | 20.3 | 10.4 | 2.2 | 12.5 | 4.1 | 1.9 | 16.3 | 4.0 | 4.3 | 4.3 |
| Human gut metagenome | 6.6 | 0.7 | 7.3 | 3.2 | 0.9 | 4.0 | 0.5 | 0.8 | 4.5 | 3.3 | 3.9 | 3.9 |
| Human RNA-seq (k=61) | 4.4 | 0.6 | 5.0 | 3.6 | 3.9 | 7.5 | 1.1 | 2.4 | 7.1 | 4.3 | 2.3 | 2.3 |

### 7.2 Evaluation of UST-Compress

We measure the compressed space-usage (Table 2), compression time and memory (Table 3), and decompression time and memory. We compare against the following lossless compression strategies: 1) the binary output of the *k*-mer counters DSK [18], KMC [17], and Squeakr-exact [20]; 2) the original FASTA sequences, with headers removed; 3) the maximal unitigs; and 4) the BOSS representation [31] (as implemented in COSMO [42]). In all cases, the stored data is additionally compressed using MFC (for nucleotide sequences, i.e. 2 and 3) or LZMA (for binary data, i.e. 1 and 4). The second strategy (which we already discussed in Section 1.1) is not a *k*-mer compression strategy per say, but it is how many users store their data in practice. The fourth strategy uses BOSS, the empirically most space efficient exact membership data structure according to a recent comparison [35]. We include this comparison to measure the advantage that can be gained by not needing to support membership queries. Note that strategies 1 and 2 retain count information, unlike strategies 3 and 4. Squeakr-exact also has an option to store only the *k*-mers, without counts.

First, we observe that compared to the compressed native output of *k*-mer counters, UST-Compress reduces the space by roughly an order-of-magnitude; this however comes at an expense of compression time. When the value of *k* is increased, this improvement becomes even higher; as *k* nearly doubles, the UST-Compress output size remains the same, however, the compressed binary files output by *k*-mer counters approximately double in size. Our results indicate that when disk space is a more limited resource than compute time, SPSS-based compression can be very beneficial. Second, we observe a 4-8x space improvement compared to just compressing the reads FASTA file. In this case, however, the extra time needed for UST compression is balanced by the extra time needed to recount the *k*-mers from the FASTA file. Therefore, if all that is used downstream are the *k*-mers and possibly their counts, then SPSS-based compression is again very beneficial. Third, UST-Compress uses between 39 and 48% less space than BOSS, with comparable construction time and memory. Fourth, compared to the other SPSS-based compression (based on maximal unitigs), UST-Compress uses 10 to 29% less space, but has 10 to 24% slower compression times (with the exception of the *k* = 61 dataset, where it compresses 6% faster). The ratio of space savings after compression closely parallels the ratio of the weights of the two SPSS representations (Table 1). Fifth, we note that the best compression ratios achieved are significantly better than the worst case Conway Bromage lower bound of *>* 35 bits per *k*-mer for the *k* = 31 datasets and 95 bits per *k*-mer for the *k* = 61 dataset. Finally, we note that the differences in the peak construction memory, and the total decompression run time and memory (*<* 2min and *<* 1 GB for UST-Compress, respectively, table not shown) were negligible.

We also compressed a subset of samples from a de-noised index of 450,000 microbial DNA data used recently in large scale indexing projects of BIGSI [43] and COBS [44]. Each sample consists of error-corrected 31-mers (without abundance information) from a corresponding sequencing experiment, natively stored as bzipped McCortex binary file (see [43, 44] for details). We downloaded 19,000 of these files from [45]. We ran UST-Compress, which reduced the disk space from 507 GB to 14.7 GB, a 35x reduction. The compression took a total of 82 hours and a peak memory of 3 GB (using one core).

### 7.3 Evaluation of UST-FM

We measure the memory taken by the data structure (Table 4), the query times (Table 5), and the time and memory taken during construction (Table S4). We compare UST-FM against two other space-efficient exact static membership data structures for *k*-mer sets. The first builds the FM index on top of the maximal unitigs (we refer to this as unitig-FM, but it referred to originally as dbgfm in [4]). The second is BOSS, which, as mentioned previously, was shown [35] to have superior space usage. We did not compare against the Bloom filter trie [36], which is fast but uses an order of magnitude more memory than BOSS [35]. Other data structures, such as Pufferfish [8], blight [9], and Bifrost [10], implement more sophisticated operations and hence use significantly more memory than BOSS. Moreover, these make use of a unitig SPSS representation and hence could potentially themselves incorporate the UST approach.

**Table 4:**
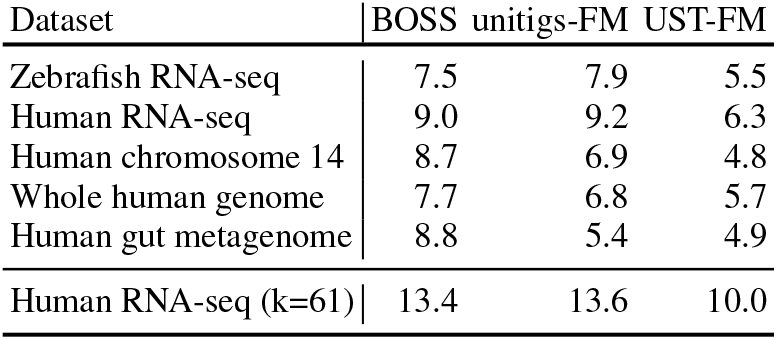
UST-FM data structure size, shown in the average number of bits per distinct *k*-mer in the dataset. This was measured by taking the peak memory usage during membership queries.

**Table 5:** UST-FM query time (in seconds) for two sets of 10,000 *k*-mers each, using the Human RNA-seq indices. The first set contains *k*-mers drawn from the dataset, so that UST-FM returns a hit. The second set takes randomly generated *k*-mers which were verified to not be present in the dataset. We measured the query times (per *k*-mer) after the index was already loaded into memory.

|  | BOSS | unitigs-FM | UST-FM |
| --- | --- | --- | --- |
| $k = 31$ | | | |
| $x \in K$ | 3.80 | 0.51 | 0.49 |
| $x \notin K$ | 1.48 | 0.38 | 0.37 |
| $k = 61$ | | | |
| $x \in K$ | 15.25 | 1.61 | 1.58 |
| $x \notin K$ | 5.10 | 0.35 | 0.37 |

First, the UST-FM index is 25 - 44% smaller and the queries are 4 to 11 times faster compared to BOSS; however, it takes 2 to 5 times longer to build. This time is dominated by FM-index construction [41], rather than by UST. Second, the UST-FM index is 10 - 32% smaller than the unitigs-FM index, with negligibly faster query time. Finally, the memory use during construction was similar for all approaches.

## 8 Conclusion

In this paper, we define the notion of a spectrum-preserving string set representation of a set of *k*-mers, give a lower bound on what could be achieved by such a representation, and give an algorithm to compute a representation that comes close to the lower bound. We demonstrate the applicability of the SPSS definition by using our algorithm to substantially improve space efficiency of the state-of-the-art in two applications.

A natural question is why we limit ourselves to SPSS representations. One can imagine alternative strategies, such as allowing a *k*-mer to appear more than once in the string set, or allowing other types of characters. In fact, for any concrete application, one might argue that a SPSS representation is too restrictive and can be improved. However, we chose to focus on SPSS representations because they are the common denominator in the applications of unitig-based representations we have observed [4,8–10]. In this way, they retain broad applicability, as opposed to more specialized representations.

One limitation of UST is the time and memory needed to run Bcalm2 as a first step. Bcalm2 works by repeatedly gluing *k*-mers into longer strings, taking care to never glue across a unitig boundary. However, this care is wasted in our case, since UST then greedily glues across unitig boundaries anyway. Therefore, a potentially significant speedup and memory reduction of UST would be to implement it as a modification of Bcalm2, as opposed to running on top of it. This can keep the high-level algorithm the same but change the implementation to work directly on the *k*-mer set by incorporating algorithmic aspects of Bcalm2.

## Acknowledgements

We are grateful to Rayan Chikhi for feedback and help with modifying Bcalm2. PM and AR were supported by NSF awards 1453527 and 1439057. AR is supported by NIH Computation, Bioinformatics, and Statistics training program.

## A Appendix

**Table S1:** Dataset Characteristics. Singletons are not included in the *k*-mer count. Unless otherwise stated, *k* = 31.

| Dataset | Source | # reads | Read length (bp) | # distinct $k$-mers |
| --- | --- | --- | --- | --- |
| Zebrafish RNA-seq | SRX3022435 | 59,741,039 | 101 | 124,740,993 |
| Human RNA-seq | SRR957915 | 49,459,840 | 101 | 101,017,526 |
| Human chromosome 14 | GAGE [46] | 36,504,800 | 101 | 99,941,572 |
| Whole human genome | SRR034939 | 36,201,642 | 100 | 391,766,120 |
| Human gut metagenome | SRR341725 | 25,479,128 | 90 | 103,814,001 |
| Human RNA-seq ($k = 61$) | SRR957915 | 49,459,840 | 101 | 75,013,109 |

**Table S2:**
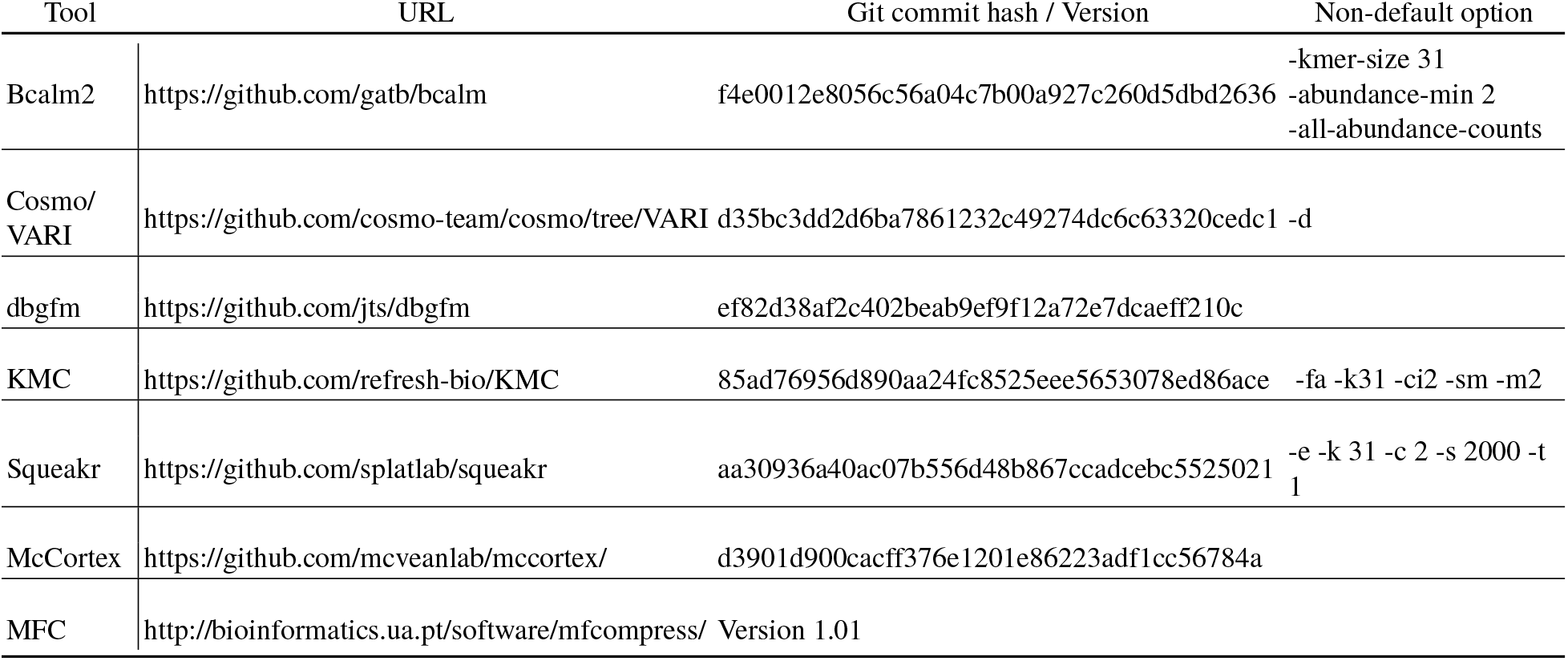
Versions of the tools used in experiments. Default options were used except as noted in the last column. We show the options for *k* = 31. Reproducibility details are available at https://github.com/medvedevgroup/UST/tree/master/experiments

**Table S3:** The percent of *cdBG*(*K*) vertex-sides that belong to isolated vertices, that are dead-sides, and that are counted by *nsp*.

| Dataset | $2n_{iso}$ | $n_{dead}$ | $n_{sp}$ |
| --- | --- | --- | --- |
| Zebrafish RNA-seq | 13 | 18 | 21 |
| Human RNA-seq | 13 | 20 | 18 |
| Human chromosome 14 | 10 | 14 | 21 |
| Whole human genome | 54 | 8 | 9 |
| Human gut metagenome | 44 | 10 | 15 |
| Human RNA-seq ($k = 61$) | 24 | 22 | 15 |

**Table S4:**
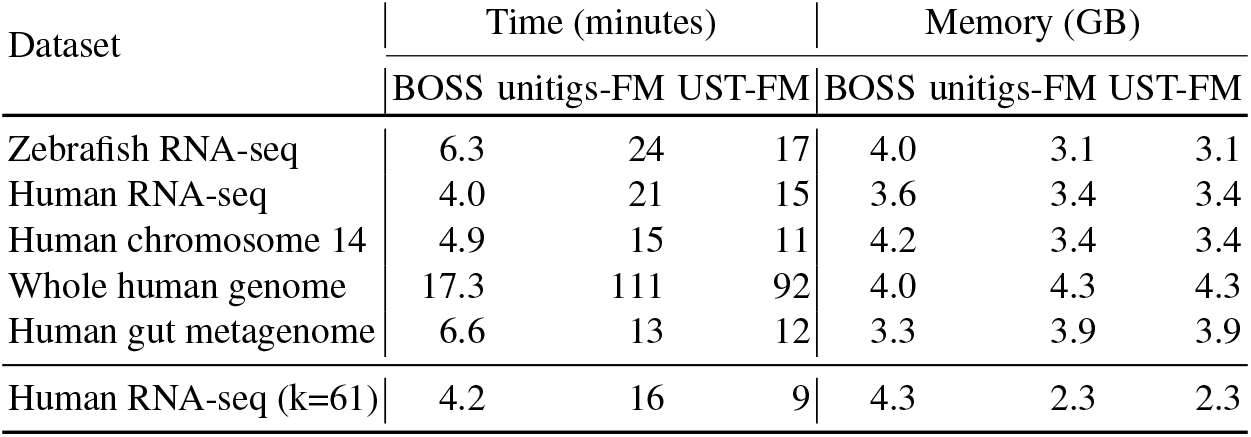
Time and memory for construction of index by UST-FM and others.

